# Exploring multiple sensory systems in ovipositors of *Drosophila suzukii* and related species with different egg-laying behaviour

**DOI:** 10.1101/651091

**Authors:** Cristina Maria Crava, Roberto Romani, Damiano Zanini, Simone Amati, Giorgia Sollai, Roberto Crnjar, Albrecht Haase, Marco Paoli, Marco Valerio Rossi-Stacconi, Omar Rota-Stabelli, Gabriella Tait, Gianfranco Anfora

## Abstract

*Drosophila suzukii* is an invasive agricultural pest species that lays eggs in fruit during ripening, while most closely related *Drosophila* species use rotten matter as oviposition substrates. This behaviour is allowed by an enlarged and serrated ovipositor that can pierce intact fruit skin. *D. suzukii* combines multiple sensory systems (mechanosensation, olfaction, and taste) to select oviposition sites. Here, we test the hypothesis that the *D. suzukii* ovipositor is involved in these sensory modalities. We first investigate the ovipositor gene expression using a comparative framework of four *Drosophila* species with gradual changes in ovipositor morphology to identify evolutionary adaptations specific to *D. suzukii*. Results show transcription of chemoreceptors and mechanoreceptors in the four species, with a common core of sensory receptors expressed in all of them. Then, we demonstrate that sensory structures present in the distal tip of the *D. suzukii* ovipositor are mechanosensory-like sensilla, and that the degenerin/epithelial sodium channel *ppk* is expressed in homologous structures in *Drosophila melanogaster*. Our results suggest the *D. suzukii* ovipositor playing a role in mechanosensation, which might be shared with other *Drosophila* species.

## BACKGROUND

*Drosophila suzukii* (Matsumura) (Diptera Drosophilidae), also called spotted wing drosophila, is an invasive South Eastern Asian fly species that was identified outside its native range in California (US) in 2008 and in Spain and Italy (Europe) in 2009 (1, 2). Since then, it has quickly spread across both continents and settled down in several countries, where it represents a major threat for soft fruit production (3). Differently from the majority of drosophilids, which thrive and lay eggs on already damaged or rotting vegetal substrates, *D. suzukii* is able to pierce and lay eggs on healthy ripening fruits before harvesting. Wherever it is present, this causes extensive agricultural damage and has boosted research on the ecology and chemosensory behaviour of this pest with the aim to find innovative, effective, and eco-friendly methods to reduce its attacks (reviewed in 4).

Several aspects of *D. suzukii* ecology and genetics have been analysed in a comparative *Drosophila* framework to identify key evolutionary innovations that allowed the transition from rotten to fresh fruit egg-laying behaviour (5–10). The major morphological shift from rotten fruit-ovipositing *Drosophila* species (like the insect model *Drosophila melanogaster*) is the presence of an unusually enlarged and serrated ovipositor, which is shared with the sister species *Drosophila subpulchrella,* and allows the wounding of the intact skin of berries (Figure 1) (7). This feature is not present in another closely related Asiatic spotted wing *Drosophila* species, *Drosophila biarmipes,* whose ovipositor shows intermediate features between *D. suzukii* and *D. melanogaster* (Figure 1).

**Figure 1.**
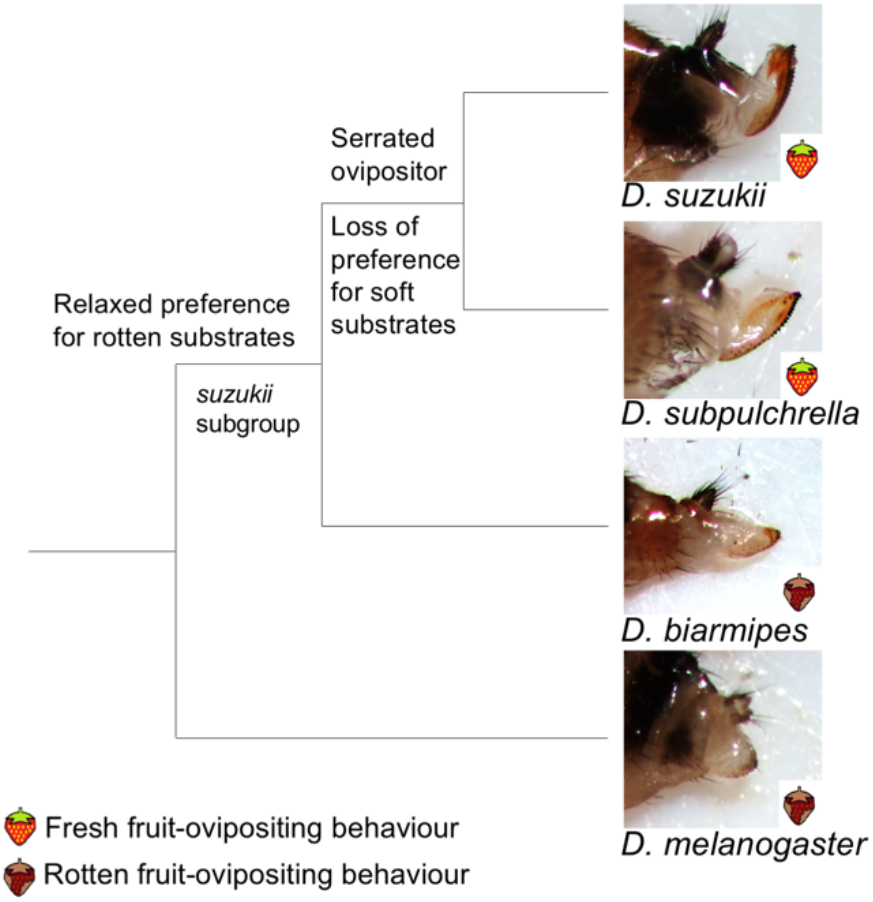
Evolutionary affinities and ovipositor shapes of *Drosophila suzukii* and other *Drosophila* species used in the study. The currently most accepted scenario for fresh fruit-egg laying behaviour evolution (9). Phylogeny is based on (7).

The ovipositor morphology correlates with the stiffness of the oviposition substrates, and serrated designs facilitate egg laying by attenuating the required penetration forces to cut through the fruit skin. In fact, *D. melanogaster* egg-laying is inhibited by stiff substrates, whereas *D. suzukii* has a broad tolerance, and *D. biarmipes* displays an intermediate behaviour (9). Besides mechanosensation, also the chemical composition of the egg-laying substrate contributes to *D. suzukii* oviposition behaviour: strawberry odours are sufficient to elicit oviposition, suggesting that olfaction has an important role in egg-laying site choice decision. Moreover, olfaction-impaired *D. suzukii* performed as wild-type flies (they prefer ripe over rotten strawberries puree) when allowed to get in touch with the oviposition substrate (9). Hence, contact chemosensory systems, such as taste, are likely used by *D. suzukii* for egg-laying site selection together with olfaction and mechanosensation. Insects detect non-volatiles compounds mainly through the gustatory receptor neurons (GRNs), which are scattered over various locations across their body. In *D. melanogaster*, GRNs house either gustatory receptors (GRs) or ionotropic receptors (IRs), and in adults they are mainly located at the labellum and along the legs (11). It has long been suggested that GRNs are also present in the ovipositor of *D. melanogaster* (12, 13) as it was demonstrated in other insect species (14–17). However, a functional characterization of these structures is lacking. An intact hydrophobic cuticle, that prevents water loss and microbial contamination, covers healthy undamaged fruits (18). We therefore conjecture that, differently from other drosophilids, which can easily taste rotten fruit substrates with legs and mouthparts, fruit cuticle of ripening fruits likely prevents *D. suzukii* probing with these appendages. The ovipositor pierces fruit skin and comes into contact with fruit flesh, thus the possibility that this organ may carry chemosensory organs which contribute to egg-laying decision is suggestive.

Here, we test this hypothesis using RNAseq to map the transcriptional profile of the female abdominal distal tip into a phylogenetical framework composed by *D. suzukii* and other three *Drosophila* species characterised by gradual changes in their ovipositor structure (from blunt to serrated ovipositor) (Figure 1). One of our aims is to understand if the presence of chemosensory-related transcripts might be a common *Drosophila* feature or something unique to fresh-fruit-ovipositing *Drosophila* species. We found expression of a set of chemosensory and mechanosensory-related mRNA conserved among the four species. This prompted a detailed analysis of the ultrastructure of pegs and sensilla present at the ovipositor of *D. suzukii*, and the use of transgenic *D. melanogaster* GAL4-drivers to provide evidence of sensory neuron-specific genes expressed in ovipositor pegs.

## METHODS

### Insects

Insects used for transcriptomics, immunohistochemistry, and electron microscopy were taken from laboratory colonies maintained at the Fondazione Edmund Mach (Italy). *Drosophila suzukii* and *D. melanogaster* strains were founded with individuals collected in 2010 in the Trento province (Italy) and periodically refreshed with insects caught from the same field sites. *Drosophila biarmipes* (genotype Dbii\wild-type, stock # 14023-0361.09) and *Drosophila subpulchrella* (Dspc\wild-type, stock # 14023-0401.00) strains were obtained from the Drosophila Species Stock Center in 2011. The four *Drosophila* species were reared on a standard diet (https://stockcenter.ucsd.edu/info/food_cornmeal.php), maintained at 23–25 °C, 65 ± 5% relative humidity and under a 16:8 h light:dark photoperiod.

*Drosophila melanogaster* transgenic strains used for imaging were obtained from the Bloomington Drosophila Stock Center and reared under the same conditions as described above. GAL4 lines are listed in Supplementary Table S1 and the reporter line expressing hexameric green fluorescent protein (6GFP) was #52262. Three to ten days old mated females were used for the analysis.

### RNA extraction and sequencing

RNA was extracted from the abdominal distal tip of 3-10 days old mated females. Dissected tissues were stored at −80 °C in RNAlater (ThermoFisher Scientific) until extraction. Each species sample was composed of RNA extracted from around 60-80 individuals. Samples were homogenized using TissueLyser (Qiagen) and total RNA was extracted with TRIzol reagent (ThermoFisher Scientific), following the manufacturer’s protocol. DNA contamination was removed with a DNase I (ThermoFisher Scientific) incubation step. A second RNA extraction with PureLink RNA Mini Kit (ThermoFisher Scientific) was performed to remove DNase and concentrate samples. Total RNA (~1 μg/sample) was sent to Beckman Coulter Genomics (Danvers, MA USA) for library preparation and Illumina sequencing. Library preparation was carried out through polyA + selection and paired-end (PE) sequencing was run on an Illumina HiSeq 2500 System with V3 chemistry that generated 100 bp reads. Raw reads are accessible from the Genbank SRA database (BioProject number PRJNA526247) (Supplementary Table S2).

### De novo *transcriptome assembly, annotation, and gene ontology*

Raw reads were trimmed with Trimmomatic (19). Both paired and unpaired reads were used for a *de novo* assembly of the transcriptome for each species with Trinity v2.0.6 (20), using the normalization step and flag --min_kmer_cov 2. The transcriptome quality was checked by mapping the paired reads against the assembled transcriptome with Bowtie (21) (Supplementary Table S2). The four transcriptomes were annotated using Standalone Blast+ (22). Blast searches were run with the command blastx using the predicted proteins from the *D. melanogaster* genome (version r6.25) as the database. The top hit for each sequence was retained when the E-value was less than 1 × 10^-10^. PANTHER version 14.0 (23) was used to extract gene ontology (GO) terms (Panther GO-Slim) for each annotated transcriptome. Venn diagrams were created using Venny 2.1.0 (24).

### Quantification of gene expression of species-specific chemosensory-related transcripts

To confirm the identity of chemosensory-related genes in each species and to estimate their expression level in each *Drosophila* species, we mapped trimmed PE reads against the dataset of manually curated annotations for *D. biarmipes, D. suzukii,* and *D. melanogaster Or, Gr, Ir,* and *Obp* genes (5, 6). For *D. subpulchrella,* we estimated expression only for *Or* and *Gr* genes using an in-house curated dataset. Proper PE trimmed reads were mapped to the annotated isoforms using Bowtie v1.0.0 (21), and the read counts were estimated by RSEM v1.2.20 (25). Genes were defined as expressed if they had at least twenty RSEM estimated expected counts. Transcripts per million reads (TPM) were visualized with heatmap.2 implemented in package gplots (R Core Team, 2015) to compare expression levels among species.

### *Reverse transcription PCR of* D. suzukii *chemosensory-related genes*

Expression of chemosensory receptor genes, found to be expressed in the *D. suzukii* abdominal distal tip by RNAseq analysis, was confirmed by reverse transcription PCR (RT-PCR). *Orco* and *Gr64,* which were not found to be expressed by RNAseq, were used as negative control and genomic DNA as positive control. The used primers are listed in Supplementary Table S3. RNA was extracted with Trizol and treated with DNAse I as described before. 1 μg RNA was then retrotranscribed to cDNA with SuperScript III Reverse Transcriptase (ThermoFisher Scientific) following the manufacturer’s protocol. To control for genomic DNA contamination, RNA underwent a parallel mock reverse transcription step in which the reverse transcriptase was omitted. Amplifications were carried out with GoTaq Green Master Mix (Promega) in a final volume of 25 μl containing 1 μl of cDNA diluted 1:10 and 0.4 μM of each primer, at the following conditions: 2 min at 95°C, then 25 cycles composed by a 30 s step at 95°C, 30 s at 55°C, and 1 min at 72°C, followed by a final elongation step of 5 min at 72°C. PCR amplicons were run on 1% agarose gel stained with Midori Green Advance (Nippon Genetics).

### Drosophila suzukii *ovipositor immunohistochemistry*

*Drosophila suzukii* adult females were anesthetized using CO_2_. Abdominal distal tips were cut with a razor blade and fixed in 4% PFA in PBS (Sigma-Aldrich) for 40 min on ice. Samples were then washed three times with PBS for 20 min, incubated in 10% sucrose (Sigma-Aldrich) solution, and kept rotating for 1 h at room temperature (RT). Sucrose was changed to 25% solution, and samples were kept rotating overnight at 4°C. Samples were then embedded in OCT (OCT mounting medium Q PATH, VWR) and mounted on a sample holder. Sections of 15 μm thickness were cut with a CM 1510-3 cryostat (Leica) and collected on a SuperFrost glass slide (ThermoFisher Scientific).

Slides were washed in PBS-T (PBS + 0.1% Triton-X-100, Sigma-Aldrich) for 5 min and then blocked in 5% normal goat serum (Sigma-Aldrich) in PBS-T for 30 min. Anti-horseradish peroxidase (HRP) cyanine-conjugated antibody (Cy3 AffiniPure Rabbit Anti-Horseradish Peroxidase, Jackson ImmunoResearch) diluted 1:300 was used to stain the neurons. Slides were kept in a moist chamber at 4°C overnight in dark. The next day, antibodies were removed, and the slides were washed three times with PBS-T for 5 min, mounted using Vectashield (Vector Laboratories), and imaged on a confocal microscope TCS SP8 (Leica).

### Drosophila suzukii *ovipositor scanning electron microscopy*

Adult females of *D. suzukii* were anaesthetized by exposure to cold temperatures (−18°C) until death, then they were immediately soaked in 60% alcohol. The ovipositor of each individual was dissected from the abdomen. Specimens were dehydrated in a series of graded ethanol, from 60% to 99%, 15 min for each step. After dehydration, 99% ethanol was substituted with pure HMDS (Hexamethyldisilazane, Sigma-Aldrich) and the specimens were allowed to dry under a hood at room temperature; this step was repeated twice. Up to five samples were mounted on aluminium stubs, with different orientations, in order to obtain a clear view on the ventral and lateral sides of the ovipositor. Mounted specimens were gold-sputtered using a Balzers Union SCD 040 unit (Balzers). The observations were carried out using a Philips XL 30 scanning electron microscope (SEM) operating at 7-10 KV, WD 9-10 mm.

### Drosophila suzukii *ovipositor transmission electron microscopy*

Ten female *D. suzukii* individuals were anesthetized by exposure to cold temperatures (−18°C) for 60 s, then immediately immersed in a solution of glutaraldehyde and PFA 2.5% in 0.1 M cacodylate buffer plus 5% sucrose, pH 7.2-7.3. The ovipositor was detached from the abdomen, reduced in size to help fixative penetration, and left at 4°C for 24 h. Then, the specimens were washed twice in cacodylate buffer for 10 min, post-fixed in 1% OsO4 for 1 h at 4°C, and rinsed in the cacodylate buffer. Dehydration in a graded ethanol series from 60% to 99%, was followed by embedding in Epon-Araldite with propylene oxide as bridging solvent. Thin sections were taken with a diamond knife on an LKB Bromma ultramicrotome (LKB) and mounted on formvar-coated 50 mesh grids. Then, sections on grids were stained with uranyl acetate (20 min, RT) and with lead citrate (5 min, RT). Finally, the sections were imaged with a Philips EM 208 transmission electron microscopy (TEM). A digital camera MegaViewIII (SIS) provided high-resolution images.

### *Examination of GAL4-driven GFP expression patterns in* D. melanogaster *ovipositor*

The native GFP signal was observed at the level of the ovipositor of females expressing the super bright 6GFP reporter under the pattern of several GAL4-drivers. Flies were anesthetized using CO2, abdominal distal tips were cut, embedded in 70% Glycerol, and imaged with the confocal microscope TCS SP8 (Leica).

## RESULTS

### Sequencing data and annotation of sensory genes

Illumina RNA-seq libraries from the abdominal distal tip of four *Drosophila* species generated an average of 60M (±2.4M SD) 100 bp paired-end reads. This resulted in four *de novo* assembled transcriptomes with contig counts ranging from 31’315 (*D. subpulchrella*) to 40’162 (*D. suzukii*) (Supplementary Table S2). On average, 70% of contigs from each of the four transcriptomes had a blastx hit against *D. melanogaster* predicted proteome (Supplementary Dataset S1). In particular, *D. suzukii* hit 11’840 unique *D. melanogaster* predicted proteins, whereas *D. subpulchrella, D. biarmipes,* and *D. melanogaster* had 10’973, 11’687, and 12’734 unique *D. melanogaster* hits, respectively. Panther functional classification revealed that the Gene Onthology (GO) composition of assembled contigs was similar among the four species (Figure 2A and Supplementary Figure S1). The most represented Molecular Function GO-slim terms were binding and catalytic activity, whereas the metabolic process, biological regulation, and cellular component organization were the most represented Biological Process GO-slim terms (Figure 2A). Of 10’774 *D. melanogaster* genes encoding for proteins that hit assembled contigs, 7294 were common among the four *Drosophila* species, thus representing the conserved transcriptional core for the distal abdominal tip (which was enriched for the ovipositor, but also included the internal genitalia, the anal plate, and some abdominal tissues). On the contrary, 131, 163, 187, and 682 genes hit specifically *D. subpulchrella, D. suzukii, D. biarmipes,* or *D. melanogaster* contigs, respectively (Figure 2B).

**Figure 2.**
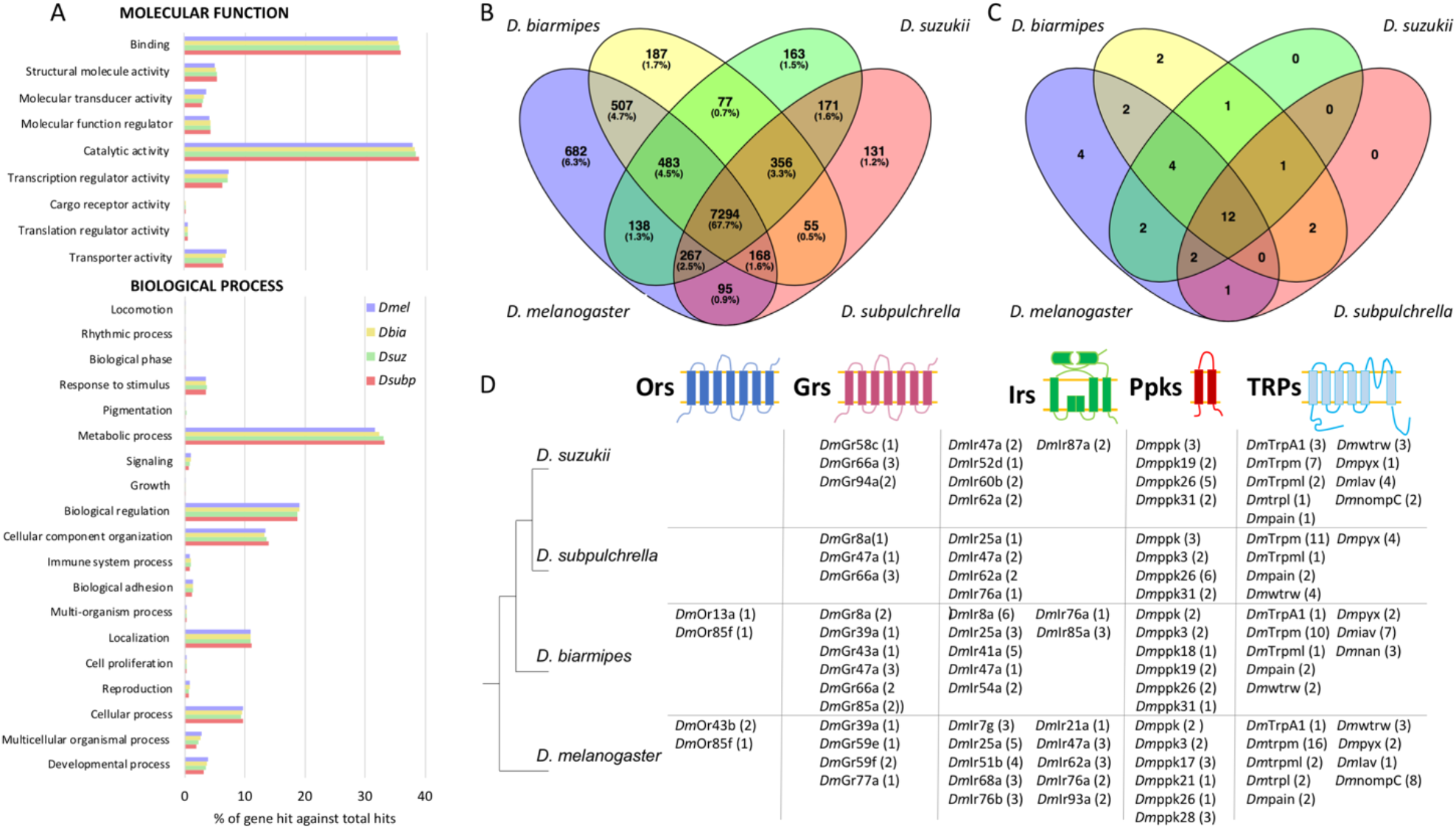
Annotation of transcriptomes from the abdominal distal tip of four *Drosophila* species reveals a core of conserved transcripts involved in sensory perception. **(A)** Gene Ontology (GO) classification for the four transcriptomes referred to Molecular Function and Biological Process (Panther GO-Slim terms). Abbreviations: Dmel, *D. melanogaster;* Dbia, *D. biarmipes*, Dsuz, *D. suzukii,* Dsubp, *D. subpulchrella*. **(B)** Venn diagram representing the unique *D. melanogaster* gene hits retrieved by blastx searches using contigs from each transcriptome as query. **(C)** Venn diagram representing the unique *D. melanogaster* odorant binding protein (Obp) gene hits retrieved by blastx searches for each transcriptome. **(D)** Schematic representation of the number of unique *D. melanogaster* gene hits encoding for sensory receptors (ORs: odorant receptors; GRs: gustatory receptors; IRs: ionotropic receptors; Ppks: pickpocket proteins; TRPs: transient receptor potential channels) retrieved in each *Drosophila* transcriptome by blastx searches. Between parentheses the number of contigs matching the same *D. melanogaster* gene hit.

Among contigs annotated by blastx searches, we various transcripts encoding for sensory receptors belonging to different classes: chemoreceptors such as odorant receptors (ORs), gustatory receptors (GRs), and ionotropic receptors (IRs), as well as other sensory receptors such as pickpocket proteins (ppks) from the degenerin/epithelial sodium channels (DEG/ENaC) gene superfamily (which contribute to mechanosensation) and transient receptor potential channels (TRPs) (which virtually contribute to every sensory modality from chemosensation to mechanosensation and thermotransduction) (Figure 2D). Of classical chemosensory receptors, 4 gustatory receptors (GRs), 4 ionotropic receptors (IRs), and 1 odorant receptor (OR) were expressed in more than one species. We also found a consistent number of contigs annotated as odorant binding proteins (OBPs) in the four transcriptomes (from 18 putative unique OBPs in *D. subpulchrella* to 27 in *D. melanogaster*). Likewise chemosensory receptors, the identity of these contigs was largely shared between the four species (Figure 2C). Of 13 *trp* genes present in *D. melanogaster* genome (26), 5 were found putatively expressed in *D. subpulchrella,* 8 in *D. biarmipes,* and 9 in *D. melanogaster* and *D. suzukii*. Of these, 5 were conserved among the four species (Figure 2D). Unique pickpocket proteins that hit assembled contigs varied from 4 (*D. suzukii* and *D. subpulchrella*) to 6 (*D. melanogaster* and *D. biarmipes*), and 3 of them were shared among all four species (Figure 2D).

### Species-specific RNAseq expression patterns of chemosensory-related transcripts

We profiled the expression levels of chemosensory-related genes in the four *Drosophila* abdominal tips by mapping their reads against available manually-annotated chemosensory gene datasets for the four species (5, 6). This led us to finely define which isoform was expressed in each species. Only genes that had ≥ 20 RSEM expected counts were considered expressed and designated as the set of abdominal distal tip genes (Supplementary Dataset S2).

We found expression of 14 chemosensory receptors in both *D. melanogaster* (3 *Grs*, 1 *Or*, and 10 *Irs*) and *D. biarmipes* (4 *Grs*, 2 *Ors*, and 8 *Irs*), and 8 in *D. suzukii* (2 *Grs*, 1 *Or*, and 5 *Irs*) (Figure 3 and Supplementary Figure S2). In *D. subpulchrella,* we tested only for *Gr* and *Or* transcription, and 2 *Grs* and 1 *Or* met our formal criteria for abdominal tip expression (Figure 3). Our analysis identified *Gr66a* and *Or43b* as expressed in the four *Drosophila* species, as well as 2 *Ir* genes were found in the three species tested for this gene family: *Ir47a* and *Ir62a*. These 4 genes likely represent a conserved core of chemoreceptors expressed in the abdominal distal tip of *Drosophila* species. We could not detect no expression of the co-receptor *Orco* in any *Drosophila* species and its absence was also confirmed in *D. suzukii* by RT-PCR (Supplementary Figure S2). Expression patterns of other chemosensory receptor genes were variable, and most of them showed a species-specific expression. Only *Gr94a* was expressed in both *D. suzukii* and *D. subpulchrella, Ir75d* in *D. suzukii* and *D. melanogaster,* and *Ir25a* in *D. melanogaster* and *D. biarmipes*.

**Figure 3.**
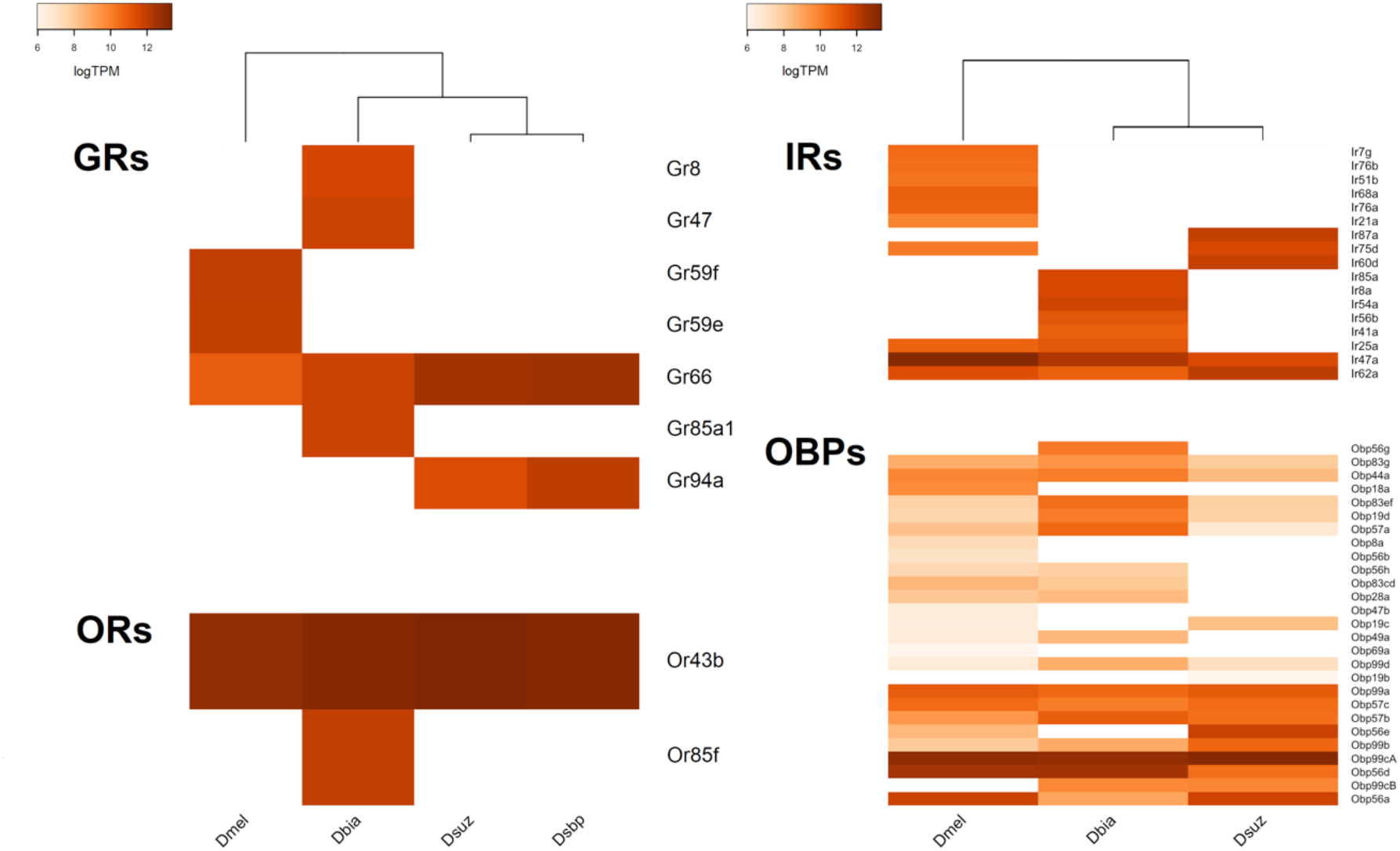
RNAseq expression levels of chemosensory-related genes in the abdominal distal tip of four *Drosophila* species. Heatmaps depict the expression level (calculated as transcripts per million, TPM) for each chemosensory-related gene found to be expressed in the most abdominal tip of *D. melanogaster* (Dmel), *D. biarmipes* (Dbia), *D. suzukii* (Dsuz), and *D. subpulchrella* (Dsbp). Expression levels were calculated using the manually curated datasets from (5, 6). For *D. subpulchrella,* we used an in-house dataset containing only GRs and ORs sequences. Abbreviations: gustatory receptors (GRs), ionotropic receptors (IRs), odorant receptors, (ORs), and odorant binding proteins (OBPs).

The dynamic range of chemosensory receptor expression in *Drosophila’s* abdominal distal tip was similar among species. Within each species, the most expressed or the second most expressed gene was *Or43b*. More generally, in each species mean expression levels of genes that shared transcription across species were substantially higher than the mean levels of the species-specific expressed genes (from 2 to 4-fold higher) (Supplementary Dataset S2).

RNAseq revealed the expression from 17 to 24 *Obp* genes in the three *Drosophila* species tested. The expression patterns of the three species largely overlapped since 13 *Obps* were expressed in *D. melanogaster, D. biarmipes,* and *D. suzukii*. On the other way around, species-specific expressed *Obps* were few: 5 in *D. melanogaster* and only 1 in both *D. biarmipes* and *D. suzukii*. Their expression levels were substantially lower than those of chemosensory receptor genes. Within each species, the mean level expression of *Obps* was from 3 to 4-fold lower than the mean level of chemoreceptors. Only *Obp99cA* had expression levels that were comparable to *Or43b* (Figure 3).

### *Ultrastructure of sensilla and pegs located on* D. suzukii *ovipositor*

Transcriptional profiles supported the hypothesis of chemosensory and mechanosensory perception in the abdominal distal tip of the four *Drosophila* species. To investigate the contribution of these sensory modalities in *D. suzukii* egg-laying behaviour, we analysed at the ultrastructural level the small bristles and pegs located at the tip of the ovipositor. Our observations revealed the presence of four different types of structures: conical pegs type 1, conical pegs type 2, trichoid sensilla and chaotic sensilla (Figure 4).

**Figure 4.**
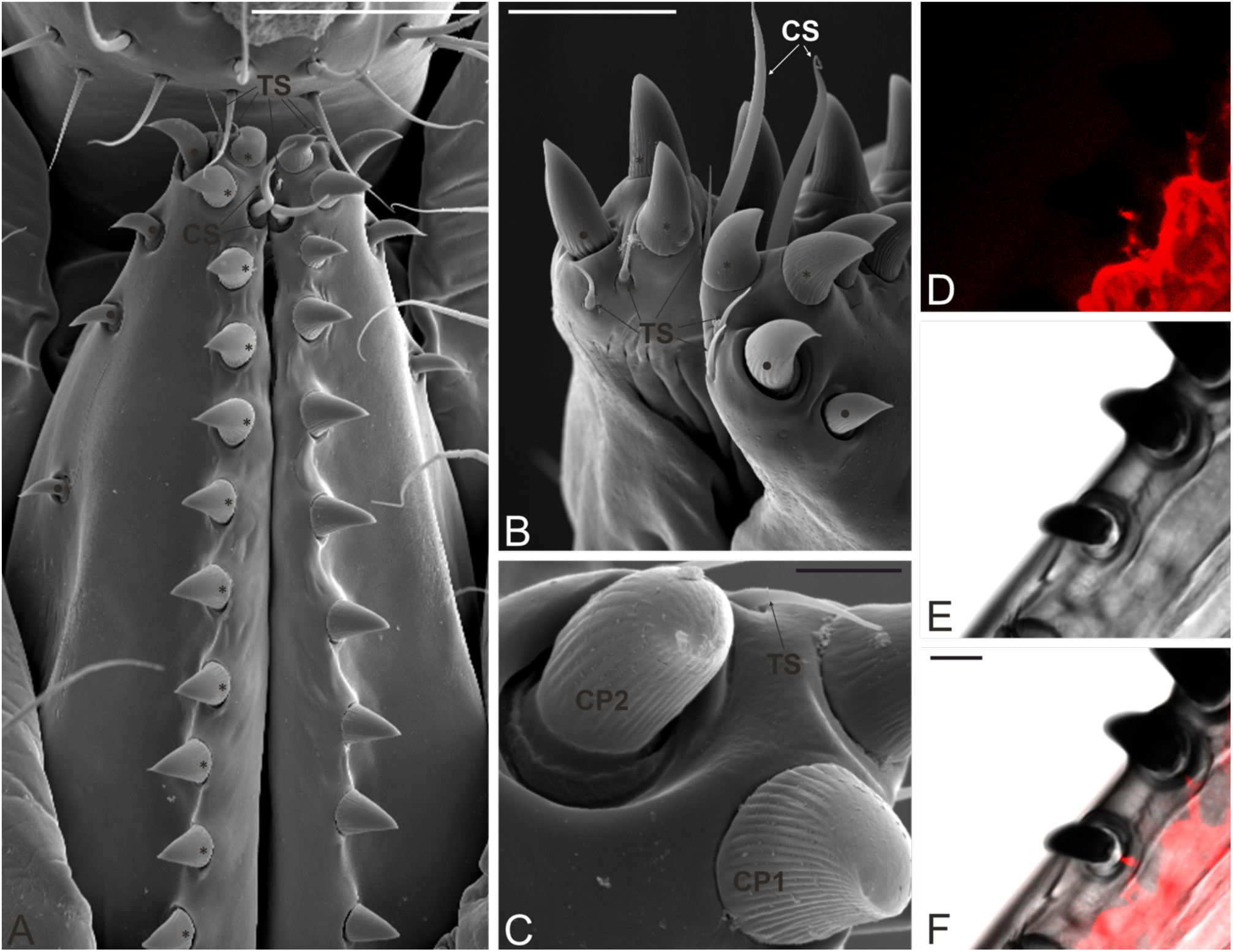
*Drosophila suzukii* ovipositor pegs and sensilla are innervated sensory structures. **(A)** Ventral view of the ovipositor of *D. suzukii* showing the two ovipositor plates and the different structures that are present. The tip of each plate presents three trichoid sensilla (TS) and a chaetic sensillum (CS). A single row of conical pegs type 1 (*) is found, with the structures arranged along the ventral edge of each ovipositor plate. Four conical pegs type 2 (•) are present, with the first one sitting at the very tip of the ovipositor plate, while the others are positioned along a medial line of the ovipositor plate. **(B)** Detailed view on the tip of the ovipositor plates. The two apical TS are clearly visible, as well as the third, inserted just behind the most apical CP1. The CS are located very close to the TS. **(C)** Close-up view of the ovipositor plate tip. The CP1 is sitting on a narrow socket; it presents a grooved cuticle that smoothens at the tip. The CP2 is sitting on a large socket; it shows a grooved cuticle as well, but with less evident grooves and a pointed tip. **(D-E)** Immunostaining of cryosection of the ovipositor plate: **(D)** neuronal anti-horseradish peroxidase signal, **(E)** bright-field, **(F)** Merged picture. Scale bars: A: 50 μm, B: 20 μm, C: 5 μm, D-F: 10 μm.

#### CONICAL PEG TYPE1 (CP1)

The outer margin of each ovipositor plate presents a single row of stout, conical pegs sitting on narrow sockets in the cuticle (Figure 4A). They are present in a number of about 15 per each plate, with a base diameter of about 10 μm and 15 μm long. CP1 are characterised by a cuticular shaft slightly bent towards the external side of the plate. The cuticle is grooved externally all along. Each structure ends in a sharp tip, although in some specimens the tip appears worn, having a blunt shape (Figure 4C and 5A). The analysis of ultrathin sections shows the internal structure of CP1, characterised by a solid, poreless cuticular shaft (Figure 5B-E). Micrographs taken at the level of the medial peg region show a thick and continuous cuticular wall with a small lumen without sensory neurons (Figure 5C). Imaging at the socket level shows as remarkable feature the presence of a single sensory neuron, typically embedded in an electron-dense dendrite sheath and ending in a tubular body (Figure 5D-E). The tubular body is located at the base of the peg, where a large socket with suspension fibres is evident (Figure 5B-D).

**Figure 5.**
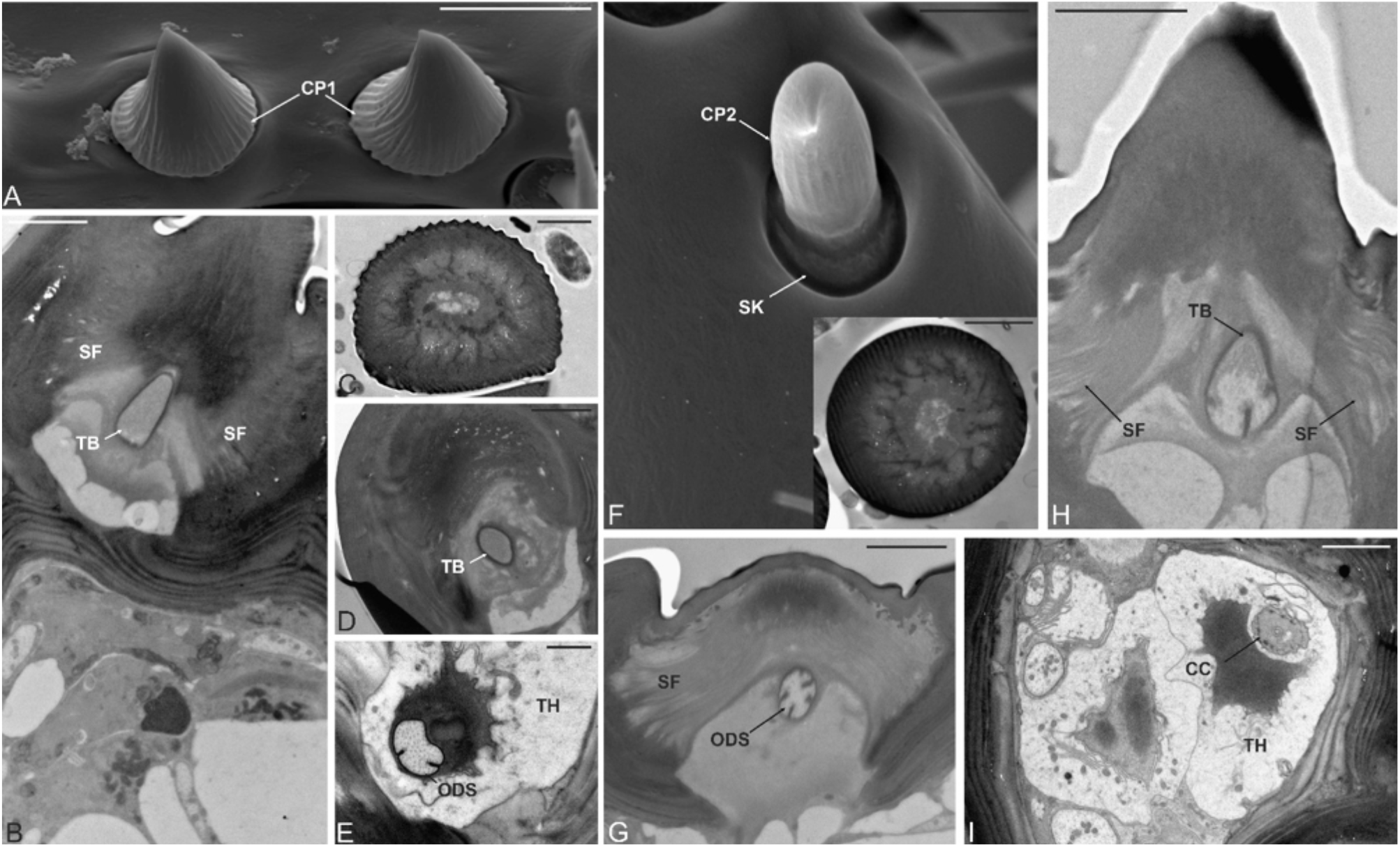
Micrographs showing details of the conical pegs type 1 and type 2 of the *Drosophila suzukii* ovipositor. **(A)** Scanning electron microscopy (SEM) ventral view of parts of the ovipositor plate ridge, showing two conical pegs type 1 (CP1). **(B)** Transmission electron microscopy (TEM) longitudinal section at the socket level. The peg is sitting on a narrow socket made of thick cuticle. Suspension fibres (SF) are apparent, holding the peg and giving flexibility to the structure. The single sensory neuron associated with the CP1 terminates in a tubular body (TB) ending just at the base of the peg. **(C-E)** Serial TEM micrographs of a CP1 cross sections, taken at different levels, show the solid cuticular structure of the peg **(C)**, the presence of the tubular body (TB) at the socket level **(D)**, and the outer dendritic segment (ODS) of the sensory neuron enclosed by the thecogen cell (TH) **(E)**. **(F)** SEM ventral view of the ovipositor plate, showing one of the conical pegs type 2 (CP2), with a slightly grooved cuticle and a sharp tip. Noticeable is also the large socket (SK) on which the CP2 is sitting in the cuticular wall of the ovipositor plate. The inset in **(F)** shows the TEM micrograph of a CP2 cross section, taken at half of the length of the peg: the peg is made of solid, thick cuticle and presents a reduced lumen, devoid of sensory neurons. **(G-I)** Serial TEM micrographs of a CP2, longitudinally and cross sections taken at different levels. They show: in **(G)** the base of a CP2 with a large flexible socket and several suspension fibres (SF). The single sensory neuron ends in a tubular body (TB) at the base of the peg. In **(H)** the large socket is visible, as well as the outer dendritic segment (ODS) of the sensory neuron. In **(I)** a cross section is imaged at a lower level respect to the previous: the sensory neuron appears at the ciliary constriction level (CC), and it is enclosed by the thecogen cell (TH). Scale bars: A: 10 μm, B-D: 2 μm, E: 1 μm, F: 5 μm, inset in F: 2 μm, G-I: 2 μm.

#### CONICAL PEG TYPE2 (CP2)

Starting from the tip of each plate, a series of 4 curved sensilla are found. They are arranged in line and show decreasing sizes from the most apical one (the larger, 14.5 μm long and 6 μm of base diameter) to the proximal one (12.5 μm long and 3.5 μm of base diameter) (Figure 4A). The cuticular shaft is slightly grooved along the longitudinal axis for most of its length, although the grooves are not as evident as in CP1 (Figure 4C). Each CP2 ends in a fine tip that can be absent in case of mechanical worn. The peg is sitting on an evident socket within the cuticular wall of the plate (Figure 5F). TEM investigation revealed also for CP2 an internal structure similar to the one described for CP1, *i.e*. the presence of a solid cuticular shaft, devoid of pores (see inset in Figure 5F), a small internal lumen without sensory neurons, and a single sensory neuron with a distal tubular body attached at the base of the peg (Figure 5G-I). The peg itself is attached flexibly to the cuticle through a large socket with an abundance of suspension fibres (Figure 5G-H).

#### TRICHOID SENSILLA (TS)

At the very tip region of each ovipositor plate, we observed 3 small TS (Figure 4A). Two of them are located on the lower side of the plate (just below and almost in between the largest CP2 and the apical CP1). The third TS is located apically on the medial region of the plate, posteriorly to the apical CP1 (Figure 4B and Supplementary Figure S3). TS are slender and finely tipped sensilla with a smooth cuticular shaft devoid of cuticular pores (12.5 μm long and 1.5 μm of base diameter). These sensilla are sitting in the cuticular wall on distinct sockets, under an angle that makes them run almost parallel to the plate cuticular wall itself. TEM images revealed that TS are made of solid cuticle, there are no pores on the cuticle and no sensory neurons entering the peg lumen (Supplementary Figure S3). A single sensory neuron was found to be associated with TS sensilla, reaching the sensillum base (Supplementary Figure S3).

#### CHAETIC SENSILLA (CS)

Each ovipositor plate shows the presence of a single CS. This sensillum is positioned apically, located medially on the ovipositor plate right behind the most apical CP1 and very close to TS. CS is long (38 μm) and slender (2.5 μm of base diameter), with a typical curved shape and a very fine tip (Figure 4B and Supplementary Figure S3). It is sitting on a large socket in the plate wall. Externally, the CS wall is smooth. Serial ultrathin sections revealed that the CS cuticular shaft is made of solid cuticle and shows a central lumen not innervated (Supplementary Figure S3). At the base, the shaft is attached to the cuticle and suspended through an elaborated socket with numerous suspension fibres (Supplementary Figure S3). A single sensory neuron was found in this area, it ends in a tubular body that attaches at the sensillum base (Supplementary Figure S3).

### *Neuronal staining of* D. suzukii *ovipositor pegs*

Anti-HRP was used as a marker to stain the neuronal membrane of the peripheral nervous system (27). HRP labelling showed the presence of a single neuron innervating each CP1s and CP2s (Figure 4D-E and Supplementary Figure S4). Neurons stopped at the CP base level, as evidenced by TEM imaging (Figure 5B and 5G). Altogether, these results suggest that the structures located on *D. suzukii* ovipositor tip host mechanosensory and not chemosensory neurons. If this inference is correct, then these neurons should not express any of the chemosensory-related genes, whose expression was found in the abdominal distal tip by RNAseq analysis, but only mechanosensory-related genes. To test this hypothesis, we moved to the model organism *D. melanogaster,* whose available transgenic toolbox allowed to drive GPF expression in its ovipositor pegs tip (Figure 1).

### *GAL4-driven GFP expression patterns of sensory genes in* D. melanogaster *ovipositor pegs*

In our systematic GAL4 expression analysis, we examined 5 genes encoding chemosensory receptors (*Or43b, Gr66a, Ir47a, Ir62a,* and *Ir75d*) and 4 mechanosensory-related genes (iav, *nompC, pain,* and ppk), which were found to be expressed by RNAseq analyses in both *D. melanogaster* and *D. suzukii* abdominal distal tip. One mechanosensory-related gene (nan) which was found to be expressed only in *D. melanogaster* and the odorant receptor co-receptor *Orco,* whose expression was not found in any transcriptome. We also examined 7 GAL4 drivers commonly used as markers for mechanosensory neurons and one pan-neuronal marker (28, 29). Using these GAL4 lines to drive 6GFP expression, we examined expression in the pegs and sensilla-like structures present on the ovipositor tip of *D. melanogaster*.

No *GAL4* drivers representing any gene encoding chemosensory receptors or *Orco* showed expression in sensory-like structure present in *D. melanogaster* ovipositor. Only *GAL4* drivers representing the pan-neuronal marker *n-syb* (Figure 6A-C) and the sodium channel *ppk* (Figure 6D-E) were expressed in the pegs, thus supporting a role in mechanosensory perception for *D. melanogaster* ovipositor pegs very similar to *D. suzukii*.

**Figure 6.**
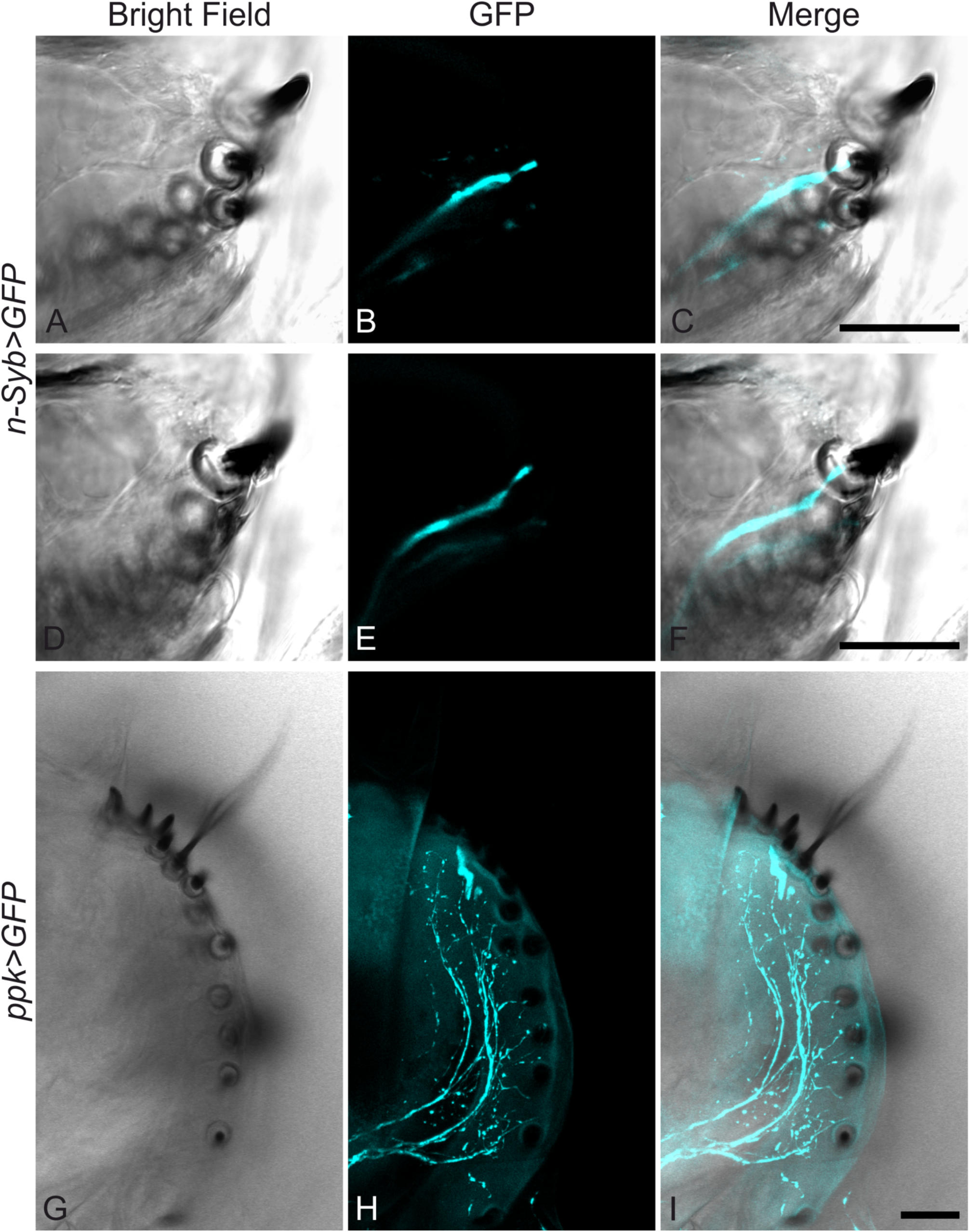
Gene expression at the *Drosophila melanogaster* ovipositor pegs. **(A-F)** The pan-neuronal marker *n-syb* showed a single neuron innervating all ovipositor pegs. **(G-I)** *ppk-GAL4* driver that labels ovipositor pegs. All scale bars 20 μm.

## DISCUSSION

Our results represent the first step toward a full molecular, anatomical, and physiological characterization of sensory perception in the *D. suzukii* ovipositor and its closely related species. In particular, these findings have (i) provided a qualitative overview of sensory genes expressed in the abdominal distal tip of four *Drosophila* species with different ovipositor shapes and egg-laying behaviours, have (ii) shown that pegs and sensilla present in the *D. suzukii* ovipositor tip have a mechanosensilla-like structure, and have (iii) identified the sodium channel *ppk* as the sensory receptor expressed in the *D. melanogaster* ovipositor pegs.

It has long been suggested that chemoreceptors are present in *D. melanogaster* female genitalia (12, 13, 30) but to our knowledge, this hypothesis has never been fully proven. Our transcriptome results have revealed that transcription of few chemoreceptors in the abdominal distal tip is a common feature in *Drosophila* species. However, both the ultrastructure of *D. suzukii* ovipositor’s pegs and sensilla, as well as the GAL4-driven GFP expression in *D. melanogaster* ovipositor’s pegs pointed out that these sensory structures host mechanoreceptors but not chemoreceptors. This suggests that the distal tip of *Drosophila* ovipositors is not used to evaluate the chemical composition of the egg-laying substrate.

On the other hand, the presence of a common transcriptional core of several homologous genes encoding for chemoreceptors suggests a conserved functional role of these proteins in the abdominal distal tip of *Drosophila* species. Of these, *Gr66a* is among the best characterized *D. melanogaster* GRs. It is normally expressed in broadly-tuned bitter-sensing neurons and its canonical function is the detection of aversive compounds (31–34). Its expression is not restricted to taste neurons (30), but it has been also identified in multidendritic neurons in the adult *Drosophila* abdomen (35), which may be the origin of the transcripts detected in our study. Other taste receptors for which we observed a transcriptional overlap among species are *Ir47a* and *Ir62a*, whose expression in *D. melanogaster* have been identified by GAL4-driven GFP expression in taste neurons in leg and labellum (36). Recently, *Ir62a* has been shown to be required together with *Ir25a* and *Ir76b* in specific GRNs to taste Ca^2+^ in food (37). Consistent with this function, in the *D. melanogaster* transcriptome we found the expression of those three transcripts. It has been shown that, besides their main function in taste, some GRs (and likely some IRs) may have other non-gustatory functions, such as the detection of internal ligands. For example, *Gr43a* is involved in fructose detection in the *D. melanogaster* brain, and it is also expressed in the female uterus, possibly to sense fructose presence in the seminal fluid (38). We did not detect expression of *Gr43a* in any of the four transcriptomes generated in this study, but the expression of the other taste receptors found may have similar physiological roles in internal sensing. Internal sensory neurons are present in the reproductive tract of *D. melanogaster* to sense sex peptide (39, 40), and this may be another location where chemoreceptor transcripts are expressed. Interestingly, in all *Drosophila* species we could detect the expression of *Or43b* homologs (which in *D. melanogaster* is normally expressed in the ab8A antennal sensilla) but not of *Orco,* whose co-expression is normally required for odour responsiveness in olfactory organs (41). As discussed before for taste receptors, *Or43b* expression localized outside its characteristic pattern may be associated with the detection of internal ligands, such as monitoring the internal nutritional state in the final gastrointestinal tract or seminal fluid evaluation.

*Drosophila suzukii* females use mechanosensation to choose the oviposition site (9, 42, 43). However, the localization of mechanosensory neurons responsible for this behaviour had not been shown yet. Here, we propose that the sensilla and pegs present on the ovipositor tip of *D. suzukii* are the sensory structures responsible to probe substrate stiffness for the egg-laying site selection. It is also possible that these sensory structures work together with other sensory organs, providing the information when the ovipositor has penetrated the substrate and peristaltic waves can start to push down the egg. Our transcriptome analysis found the expression of several genes encoding for receptors known to be expressed in proprioceptive neurons and nociceptors (44–49). The expression of most of these transcripts seems to be conserved among the four *Drosophila* species suggesting a conserved functional role. These genes represent the best candidates to be expressed in the ovipositor pegs and sensilla of *D. suzukii*.

Mechanosensation is important also for *D. biarmipes* and *D. melanogaster* egg-laying site choice (9), and we hypothesized that pegs and sensilla present in the tip of all *Drosophila* ovipositors may host mechanosensory-like neurons similar to those observed in *D. suzukii*. Hence, we used GAL4-driven GFP expression to identify genes expressed in *D. melanogaster* ovipositor pegs. Expression of *nsyb* showed that these appendages are innervated by a neuron that stopped at the base of the peg, likewise in *D. suzukii,* thus supporting the presence of mechanosensory neurons. We also showed that these neurons express *ppk,* a mechanosensitive channel in proprioceptive neurons (45).

Taken together, these findings represent an important step in delineating the function of ovipositor pegs and sensilla in *Drosophila* species with different egg-laying behaviour. Our work builds a necessary starting point for functional studies, to assess the relationship between the structure and the function of the ovipositor sensilla, and to better understand how the egg-laying behaviour diverged across *Drosophila* species. Experiments aimed to further elucidate the mechanosensory system in the ovipositor of *D. suzukii* might allow for a development of mechanosensory-based control strategies.

