## Supplemental tables for "Exploring multiple sensory systems in ovipositors of *Drosophila suzukii* and related species with different egg-laying behaviour"

Supplementary Table S1: List of GAL4 lines.

| **Genotype** | **Purpose** | **Source** | **BDSC ID (if needed)** |
| --- | --- | --- | --- |
| w[1118]; P{y[+t7.7] w[+mC]=GMR57C10-GAL4}attP2 | Pan-neuronal | BDSC | 39171 |
| w[*]; P{w[+mC]=Or43b-GAL4.C}110t8.1 | Or43b | BDSC | 23894 |
| w[*]; P{w[+mC]=Or43b-GAL4.C}110t6.3 | Or43b | BDSC | 23895 |
| w[*]; P{w[+mC]=Orco-GAL4.W}11.17; TM2/TM6B, Tb[1] | Orco | BDSC | 26818 |
| w[*]; P{w[+mC]=Gr66a-GAL4.D}2; TI{w[+mW.hs]=GAL4}Gr93a[3] | Gr66a | 28801 | 28801 |
| w[*]; P{w[+mC]=UAS-mCD8::GFP.L}LL5/CyO; P{w[+mC]=Ir47a-GAL4.K}1/TM6B, Tb[1] | Ir47a | BDSC | 60695 |
| w[*]; Bl[1]/CyO; P{w[+mC]=Ir75d-GAL4.1995}229.1/TM6B, Tb[1] | Ir75d | BDSC | 41729 |
| w[*]; P{y[+t7.7] w[+mC]=Ir62a-GAL4.K}attP40/CyO; P{w[+mC]=UAS-mCD8::GFP.L}LL6 | Ir62a | BDSC | 60713 |
| w[*]; P{w[+mC]=iav-GAL4.K}3 | iav | BDSC | 52273 |
| y[1] w[*]; PBac{y[+mDint2] w[+mC]=iav-GAL4.P}VK00014; Df(3L)Ly, sens[Ly-1]/TM6C, Sb[1] Tb[1] | Iav | BDSC | 36360 |
| w[*]; P{w[+mC]=nan-GAL4.K}2 | nan | BDSC | 24903 |
| y[1] w[*]; wg[Sp-1]/CyO, P{Wee-P.ph0}Bacc[Wee-P20]; P{y[+t7.7] w[+mC]=nompC-GAL4.P}attP154 | nompC | BDSC | 36369 |
| w[1118]; P{y[+t7.7] w[+mC]=GMR21B03-GAL4}attP2 | pain | BDSC | 49294 |
| w[*]; P{w[+mC]=ppk-GAL4.G}2 | ppk | BDSC | 32078 |
| w[1118]; P{y[+t7.7] w[+mC]=GMR34E03-GAL4}attP2/TM3, Sb[1] | Mechanosensory neurons innervating chemosensory bristles | BDSC | 48123 |
| w[1118]; P{y[+t7.7] w[+mC]=GMR81C12-GAL4}attP2/TM3, Sb[1] | Mechanosensory neurons innervating chemosensory bristles | BDSC | 48364 |
| w[1118]; P{y[+t7.7] w[+mC]=GMR12G12-GAL4}attP2 | Mechanosensory neurons innervating chemosensory bristles | BDSC | 48527 |
| w[1118]; P{y[+t7.7] w[+mC]=GMR38B08-GAL4}attP2 | Mechanosensory neurons innervating chemosensory bristles | BDSC | 49541 |
| w[1118]; P{y[+t7.7] w[+mC]=GMR39A11-GAL4}attP2 | Mechanosensory neurons innervating chemosensory bristles | BDSC | 50034 |
| w[1118]; P{y[+t7.7] w[+mC]=GMR47C08-GAL4}attP2 | Mechanosensory neurons innervating chemosensory bristles | BDSC | 50298 |
| w[1118]; P{y[+t7.7] w[+mC]=GMR49C06-GAL4}attP2 | Mechanosensory neurons innervating chemosensory bristles | BDSC | 50417 |

Supplementary Table S2: Sequencing, *de novo* assembly and annotation statistics

|  | *D. suzukii* | *D. subpulchrella* | *D. biarmipes* | *D. melanogaster* |
| --- | --- | --- | --- | --- |
| Raw reads | 63655282 | 58076113 | 59146044 | 59497395 |
| SRA ncbi accession number | SRR8705028 | SRR8705027 | SRR8705030 | SRR8705029 |
| Paired reads after trimming | 61569501 | 56300986 | 56855248 | 57548120 |
| Assembled Trinity isoforms (contigs) | 40162 | 31315 | 32245 | 35812 |
| TSA ncbi accession number | GHHJ00000000 | GHHO00000000 | GHHN00000000 | GHHI00000000 |
| N50 | 2150 | 2315 | 3081 | 3367 |
| Bowtie mapped reads % | 76.6 | 74.2 | 84.4 | 86 |
| Contigs with at least 1 FPKM (%) | 36395 (90.6) | 28478 (90.9) | 28660 (88.9) | 31528 (88.0) |
| Contigs with a hit against *D. melanogaster* proteome (%) | 28893(72) | 22071(70.5) | 22254(69) | 24737 (69) |

Supplementary Table S3: List of primers used in *D. suzukii* RT-PCR

| Target | Left primer | Right primer |
| --- | --- | --- |
| *or43b* | CGCCTATCTGCTTGAAACGC | CTGAAGAACTGTCCGCTGGT |
| *gr66a* | ATGGCTACCGACCAGTTGAC | CGACCAATTGTGCTGTTGTC |
| *gr94a* | TCGGAATTACAGTGGCCTTC | GCGAGATGAGCAGAAAAACC |
| *gr64a* | CCGTCGAGTACAATTGGCTG | GGGACTGTAGGGAAGCAACT |
| *orco* | CACCATGACAACCTCGATGCAGCC | TTACTTGAGCTGCACCAGCAC |
| *ir47a1* | CAACTTGGATCTGGTCAGCG | CCTTGATGGTCCCTGAGGTT |
| *ir62a* | GATCCGCATCACCTGTTTCC | CGGCGAGAATGAGAAAGGTG |
| *ir60d* | AACGCCTTCATCCGGAACTA | CGAATTTGATCCCAGCCCTG |
| *ir75d* | TATTCTCCCCACACACCCAC | CGCAGCCAAAGATTCCCAAA |
| *ir87a* | TTCGCAACAGTGAGGAGGAT | ACCAATACGACCAGGGAACA |
