## Supplemental figures for "Exploring multiple sensory systems in ovipositors of *Drosophila suzukii* and related species with different egg-laying behaviour"

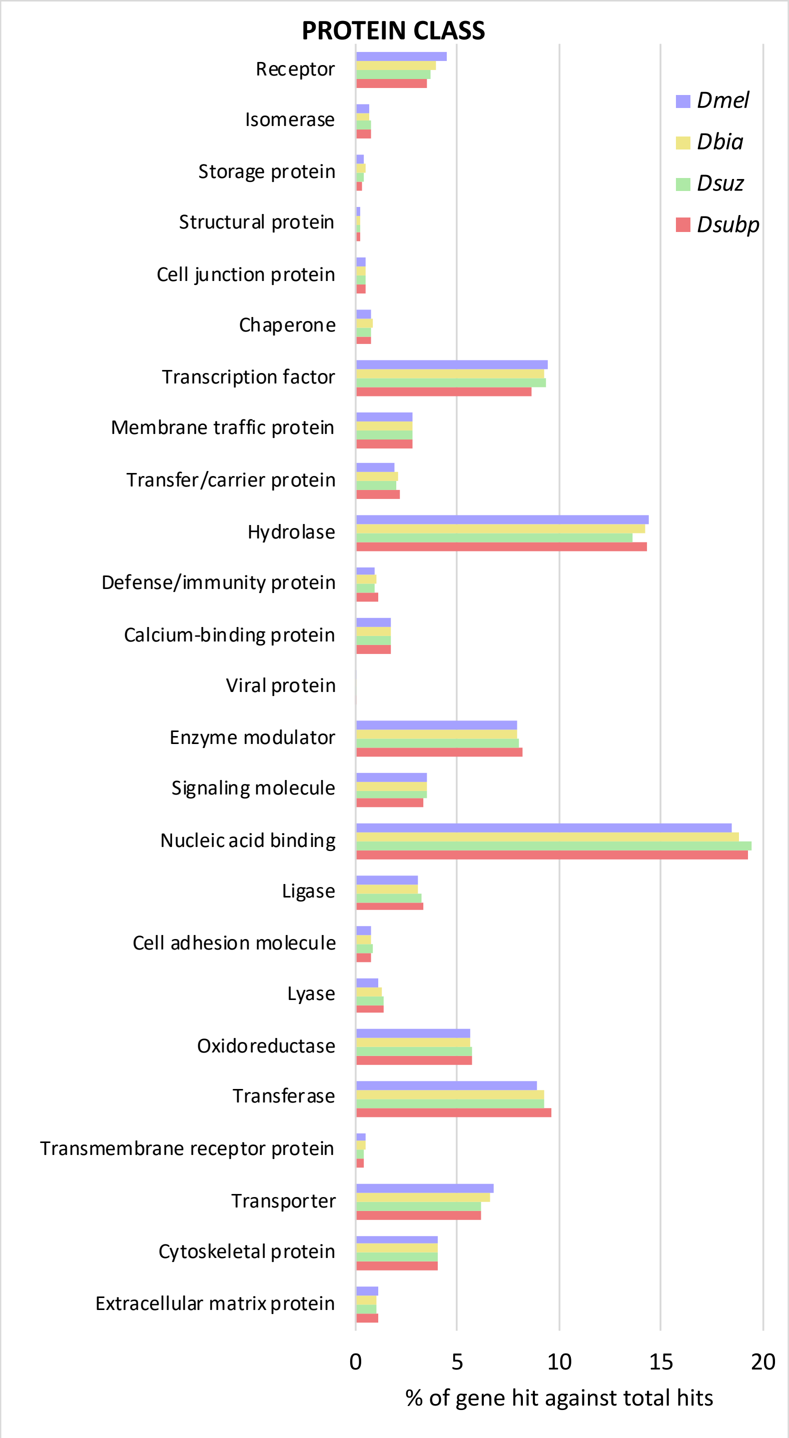


**Figure S1. Annotation of transcriptome from the abdominal distal tip of four *Drosophila* species reveals a broad core of conserved transcripts.**
Gene Ontology (GO) classification for the four transcriptomes referred to Protein Class (Panther GO-Slim terms). Abbreviations: Dmel, *D. melanogaster*; Dbia, *D. biarmipes*, Dsuz, *D. suzukii,* Dsubp, *D. subpulchrella*.

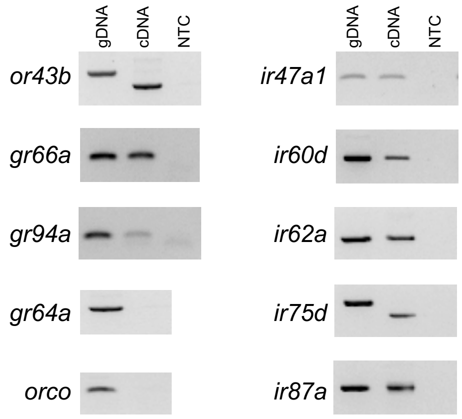


**Figure S2 Reverse transcription PCR confirmation of expression of transcripts encoding chemosensory receptors in *Drosophila suzukii* abdominal distal tip.**

Abbreviations: genomic DNA (gDNA, positive control), non-template control (NTC, negative control).

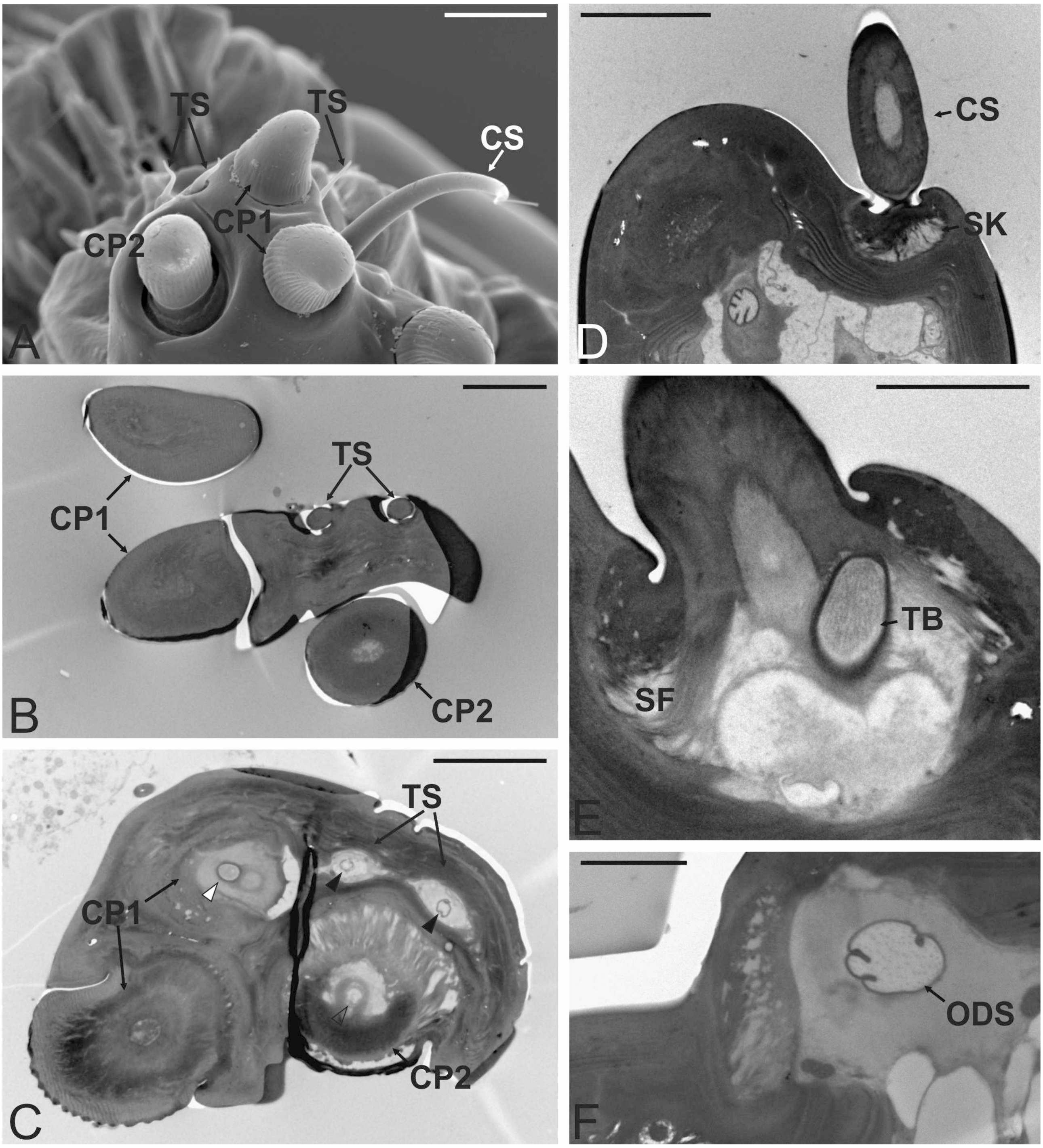


**Figure S3. Micrographs showing details of the trichoid sensilla and chaetic sensilla.**
**(A)** Scanning electron microscopy (SEM) picture of the ovipositor plate tip showing the three trichoid sensilla (TS) and the single chaotic sensilla (CS). **(B-C)** Serial transmission electron microscopy (TEM) cross sections of the ovipositor plate. In **(B)** the section is taken most apically and shows the two TS made of solid cuticle entering the cuticular wall, no sensory neurons were detected at this level. In **(C)** the two TS are pictured more proximally, the peg is no longer visible but two sensory neurons (one per each TS) are visible (black arrowheads). White and blank arrowheads show the sensory neurons associated with conical pegs type 1 (CP1) and type 2 (CP2), respectively**. (D-F)** Serial TEM longitudinal sections showing the main ultrastructural features of a CS: in **(D)** the CS is taken at the socket level (SK) and shows a thick cuticle with a central lumen without sensory neurons. In **(E)** a single sensory neuron ending in a tubular body (TB) inserted at the CS base is visible. The socket presents numerous suspension fibres (SF). In **(F)** the outer

dendritic segment (ODS) of the sensory neuron is visible. Bar scale: A: 10 µm, B-C-D: 5 µm, E-F: 2 µm.

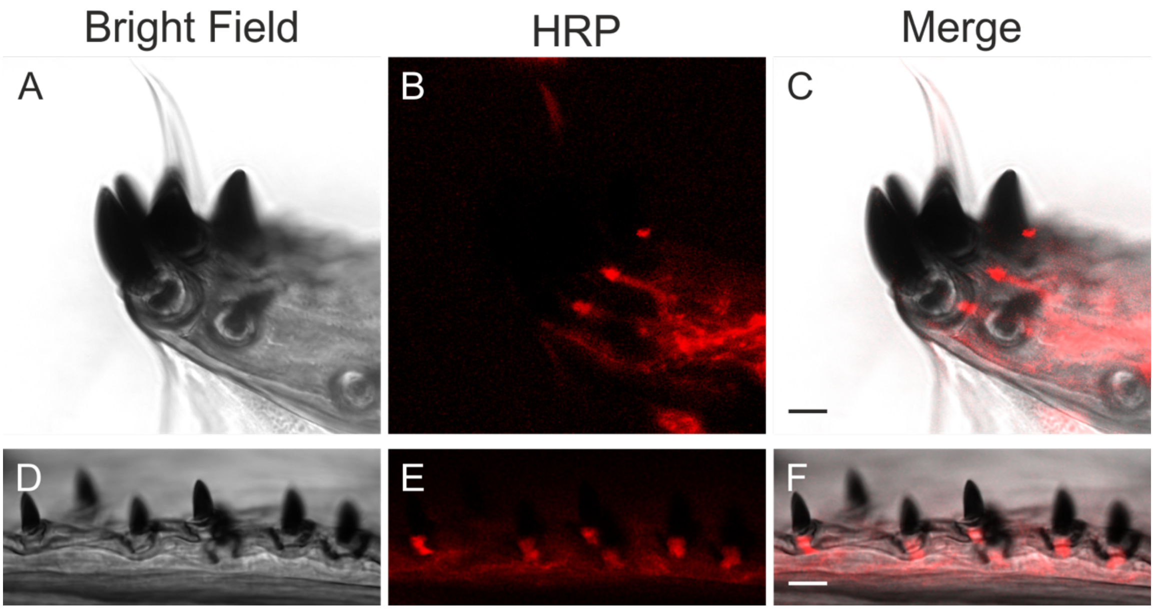

**Figure S4 Cryosection of the *Drosophila suzukii* ovipositor stained with antibody anti-horseradish peroxidase.**

Cryosection of the ovipositor plate **(A and D)** bright-field, **(B and E)** counter staining with anti-HRP to make the neuron visible, **(C and F)** Merged picture.
